# Multivariate Pattern Classification of Primary Insomnia Using Three Types of Functional Connectivity Features

**DOI:** 10.1101/532127

**Authors:** Chao Li, Yuanqi Mai, Mengshi Dong, Yi Yin, Kelei Hua, Shishun Fu, Yunfan Wu, Guihua Jiang

## Abstract

**Objective:** To investigate whether or not functional connectivity (FC) could be used as a potential biomarker for classification of primary insomnia (PI) at the individual level by using multivariate pattern analysis (MVPA).

**Methods:** Thirty-eight drug-naive patients with PI and 44 healthy controls (HC) underwent resting-state functional MR imaging. Three commonly used FC metrics (voxel-wise functional connectivity strength, large-scale functional connectivity and regional homogeneity) were calculated for each participant. We used the MVPA framework using linear support vector machine (SVM) with the three types of metrics as features separately. Subsequently, an unbiased N-fold cross-validation strategy was used to generate a classification system and was then used to evaluate its classification performances. Finally, FC metrics with significant high classification performance were compared between the two groups and were correlated with clinical characteristics, i.e., Insomnia Severity Index (ISI), Pittsburgh Sleep Quality Index (PSQI), Self-rating Anxiety Scale (SAS), Self-rating Depression Scale (SDS).

**Results:** The best classifier could reach up to an accuracy of 81.5%, with sensitivity of 84.9%, specificity of 79.1% and area under the receiver operating characteristic curve (AUC) of 83.0% (all P < 0.001). Right fronto-insular cortex, left precuneus and left middle frontal gyrus showed high classification weights. In addition, right fronto-insular cortex and left middle frontal gyrus were the overlapping regions between MVPA and group comparison. Correlation analysis showed that functional connectivity strength (FCS) in left middle frontal gyrus and head of right caudate nucleus were correlated with PSQI and SDS respectively.

**Conclusion:** The current study suggests abnormal FCS might serve as a potential neuromarkers for PI.

**Key Points:** FCS in fronto-insular cortex and middle frontal gyrus may be a neuroimaging biomarker for insomnia.

FCS can be used to distinguish between patients with primary insomnia from healthy controls with high classification accuracy (81.5%; P < 0.001).

FCS in left middle frontal gyrus and head of right caudate nucleus were correlated with PSQI and SDS respectively.

## Introduction

Primary insomnia (PI) is the most common sleep disorder and is a major risk factor for depression and in certain instances could increase mortality ^1^. At present, diagnosis for insomnia is mainly based on self-reported sleep difficulties. Objective neurobiological markers remain largely unclear and hence prevented the development of more cost-effective, efficient and accessible therapies ^2^.

Neuroimaging studies for insomnia have made substantial effort to understand the neuromechanisms of insomnia. Previous studies found aberrant brain metabolism and connectivity related to the prefrontal cortex, insular cortex, amygdala, precuneus and caudate in primary insomnia ^3-16^. For example, using PET, Nofzinger et al. ^3^ found smaller decrease in relative metabolism from waking to non-REM sleep states in the ascending reticular activating system, hypothalamus, thalamus, insular cortex, amygdala, hippocampus, anterior cingulate and medial prefrontal cortices, which supports the CNS hyperarousal hypothesis. Using independent component analysis, Michael C. Chen et al. ^6^ demonstrated that the anterior insular cortex had greater involvement with the salience network in PI. This greater involvement was also correlated with self-reported alertness and negative affect. This study highlights the importance of the salience network in hyperarousal and affective symptoms in insomnia. Diederick Stoffers et al. ^13^ found that hyper-arousal was associated with reduced caudate recruitment when performing an executive task. Interestingly, attenuated caudate recruitment did not recover after successful treatment, suggesting abnormal caudate activation is a potential vulnerability biomarker for insomnia. Recently, Lee et al. ^5^ observed that subcortical FC was changed after cognitive–behavioral therapy, which suggested that FC may be a biomarker for tracking response to treatment.

While these studies were valuable in finding relevant neuroimaging biomarkers, the studies were based on group comparisons, and hence was not sufficient for possible translational applications, such as for direct clinical diagnostic and prognostic evaluation ^17^. Up to now, it is still unclear whether or not neuroimaging could be used as a biomarker for the diagnosis of PI patients at the individual level.

In the present study, we explored whether or not three commonly used functional connectivity (FC) methods (i.e., voxel-wise functional connectivity strength, large-scale functional connectivity and regional homogeneity; please see the next section for details) could be used as potential biomarkers for the classification of individual patients with PI. This was performed using multivariate pattern analysis (MVPA) with linear support vector machine (SVM) ^18^.

## Methods

### Participants

This prospective study was approved by the ethics committee of Guangdong Second Provincial General Hospital and all participants provided written informed consent after they were provided a complete description of the study. Thirty-eight patients with PI (16 men; mean ± standard deviation age, 40.61 years ± 9.43) were recruited from the Guangdong Second Provincial General Hospital.

The inclusion criteria for PI patients were: (a) all patients must meet the Diagnostic and Statistical Manual of Mental Disorders, Fourth Edition (DSM-IV) for diagnosis of PI; (b) patients complained of difficulty falling asleep, maintaining sleep or early awakening from sleep for at least one month; (c) patients had no other sleep disorders such as hypersomnia, parasomnia, sleep-related movement disorders or other psychiatric disorders; (d) patients were younger than 60 years old (e) free from any psychoactive medication for at least 2 weeks prior to and during the study; (f) patients were right-hand dominant as assessed using the Edinburgh Handedness Inventory. Exclusion criteria were as follows: (a) Patients had an abnormal signal in any region of the brain which was verified by conventional T1-weighted or T2-fluid-attenuated inversion recovery MR imaging; (b) the insomnia disorder was caused by organic disease or severe mental disease that was secondary to depression or generalized anxiety; (c) other sleep disorders; (d) women who were pregnant, nursing, or menstruating. A total of 44 age-, gender- and education-matched healthy control subjects were recruited (11 men and 33 women; age, 39.91 years ± 9.43) from the local community by advertisements. HC met the following criterion: (a) Insomnia Severity Index score less than 7; (b) no history of swing shifts, shift work, or sleep complaints; (c) no medication or substance abuse such as caffeine, nicotine, or alcohol for at least 2 weeks prior to and during the study; (d) no brain lesions or prior substantial head trauma, which was verified by conventional T1-weighted or T2-fluid-attenuated inversion recovery MR imaging; (e) no history of psychiatric or neurological diseases; (f) right-hand dominant. All the patients were part of previous studies ^19-21^. All previous studies were investigations of between-group differences using resting-state functional MR imaging, whereas the present study explored whether resting-state functional MR imaging could be used as a neuroimaging biomarker to identify primary insomnia.

Several questionnaires were completed by the study participants. These questionnaires included the Insomnia Severity Index, the Pittsburgh Sleep Quality Index, the Self-rating Anxiety Scale, and the Self-rating Depression Scale.

### Image Acquisition

Functional MR imaging was acquired using a 1.5 Tesla MR scanner (Achieva Nova-Dual; Philips, Best, the Netherlands) in the Department of Medical Imaging, Guangdong Second Provincial General Hospital. Participants were instructed to rest with their eyes closed and remain still without falling asleep. Functional MR images were acquired in about 10 minutes using a gradient-echo planar imaging sequence as follows: interleaved scanning, repetition time/echo time = 2500ms/50 ms, section thickness = 4 mm, intersection gap = 0.8 mm, matrix = 64 × 64, field of view = 224 mm × 224 mm, flip angle = 90°, 27 axial slices, and 240 volumes. After scan, all subjects were asked if they were asleep during the scan. Those subjects fallen asleep were excluded.

### Data Preprocessing

Functional images were preprocessed using the SPM12 software package and the Data Processing Assistant for Resting-State fMRI software (DPARSF, Advanced Edition, V4.3) (http://www.rfmri.org/DPARSF) ^22^. The first 10 images of each participant were discarded to allow the signal to reach equilibrium. Subsequently, the resting-state fMRI data was corrected for temporal differences between slices and head motion. All participants had no more than 2.0 mm of maximal displacement and 2.0 of maximal rotation in any direction. Next, the corrected fMRI data were spatially normalized to the standard Montreal Neurological Institute (MNI) template, and were resampled to 3×3×3mm^3^. We further processed the data to remove linear trends and filtered temporally (band-pass, 0.01–0.1Hz). Finally, nuisance signals including 24 head motion parameters, CSF signals, white-matter signal, and global signal were regressed out from the fMRI data.

### Whole-Brain Voxel-wise Functional Connectivity Strength Analysis

Whole-brain voxel-wise functional connectivity strength as well as large-scale functional connectivity (large-scale FC) and regional homogeneity (ReHo) analysis were performed using DPARSF (http://www.rfmri.org/DPARSF). For each participant, all voxels’ time series were extracted, and then the Pearson’s correlation coefficients between the time series of all pairs of voxels were obtained to form a whole-brain voxel-wise functional connectivity matrix. Then, for each voxel, functional connectivity strength (FCS) value was calculated as the sum of the Pearson’s correlation coefficients between each voxel and all other voxels. We set a threshold of r = 0.25 to remove weak correlations. Consequently we obtained a 3D FCS map for each participant. Finally, the FCS map was converted to z scores using Fisher r-to-z transformation and further spatially smoothed with a 6 mm full-width at half maximum isotropic Gaussian kernel. It is worth noting that this computation was constrained within a gray matter mask, which was created by setting a threshold of 0.2 on the SPM12’s gray matter probability template.

### Whole-Brain Large-scale FC Analysis

Nodes were demarcated by a 268-node functional atlas ^23^, which was defined using a group-wise spectral clustering algorithm ^24^ and consequent analysis were similar to previous studies ^25^. Time series for each node was extracted for each participant by averaging the time series throughout all voxels for each node. Functional connectivity between each pair of nodes was calculated using Pearson’s correlation analysis, which produced (268×267)/2=35778 dimensional functional connectivity feature vector for each participant. Finally, Fisher r-to-z transformation was performed for functional connectivity.

### Whole-Brain ReHo Analysis

ReHo calculation was also constrained within the same gray matter mask similar to the whole-brain voxel-wise functional connectivity analysis. For each voxel of each participant, ReHo value was calculated by calculating Kendall’s Coefficient of Concordance (KCC) of the time series for the given voxel with those of its 26 neighbors ^26^. A 3D ReHo map was obtained for each participant. We further normalize the ReHo map by dividing the ReHo value for each voxel by the averaged ReHo value of the whole brain. Finally, all ReHo maps were smoothed using a 6 mm full-width at half maximum isotropic Gaussian kernel.

### Multivariate Pattern Classification Analysis

A flow chart overview of the MVPA is illustrated in Figure 1. The MATLAB codes used in our analysis are available online: https://github.com/lichao312214129/lc_rsfmri_tools_matlab/tree/master/Machine_Learning/Classification (SVM_LC_Kfold_PCA _*.m). Our analysis consisted of a 5-fold cross-validation procedure for each of the 3 modalities (i.e., FCS, large-scale FC and ReHo). At each fold k (k = 1, 2, 3, 4, 5), data of both PI and HC were divided into 2 subsets of 8 to 2. Then the 2 larger subsets from both group were fused together to form the training data (80%), with the others being test subsets (20%) and only used to assess generalization performance. Normalization and principal component dimensionality reduction were further performed on the training data. Testing data was also processed by these 2 processes using the same parameters (e.g., principal component coefficients) from the training data. We retained all the principal components to balance information loss and computational complexity of the classifier ^27^. Then, a linear SVM classifier was trained on the training data and used to classify the testing data. By comparing the predicted labels with the real labels, we acquired the classification performances (i.e., accuracy, sensitivity, specificity and area under ROC curve (AUC)) of one fold. Moreover, discriminative weights were obtained as linear SVM weights (i.e., Beta values of features from the linear SVM classifier). Final classification performances and discriminative maps were acquired as the average over the 5 folds. At the end of the iteration, we acquired the prediction labels for every participant, which was used to build the confusion matrix (please see Figure 2.A-C).

**Figure 1.**
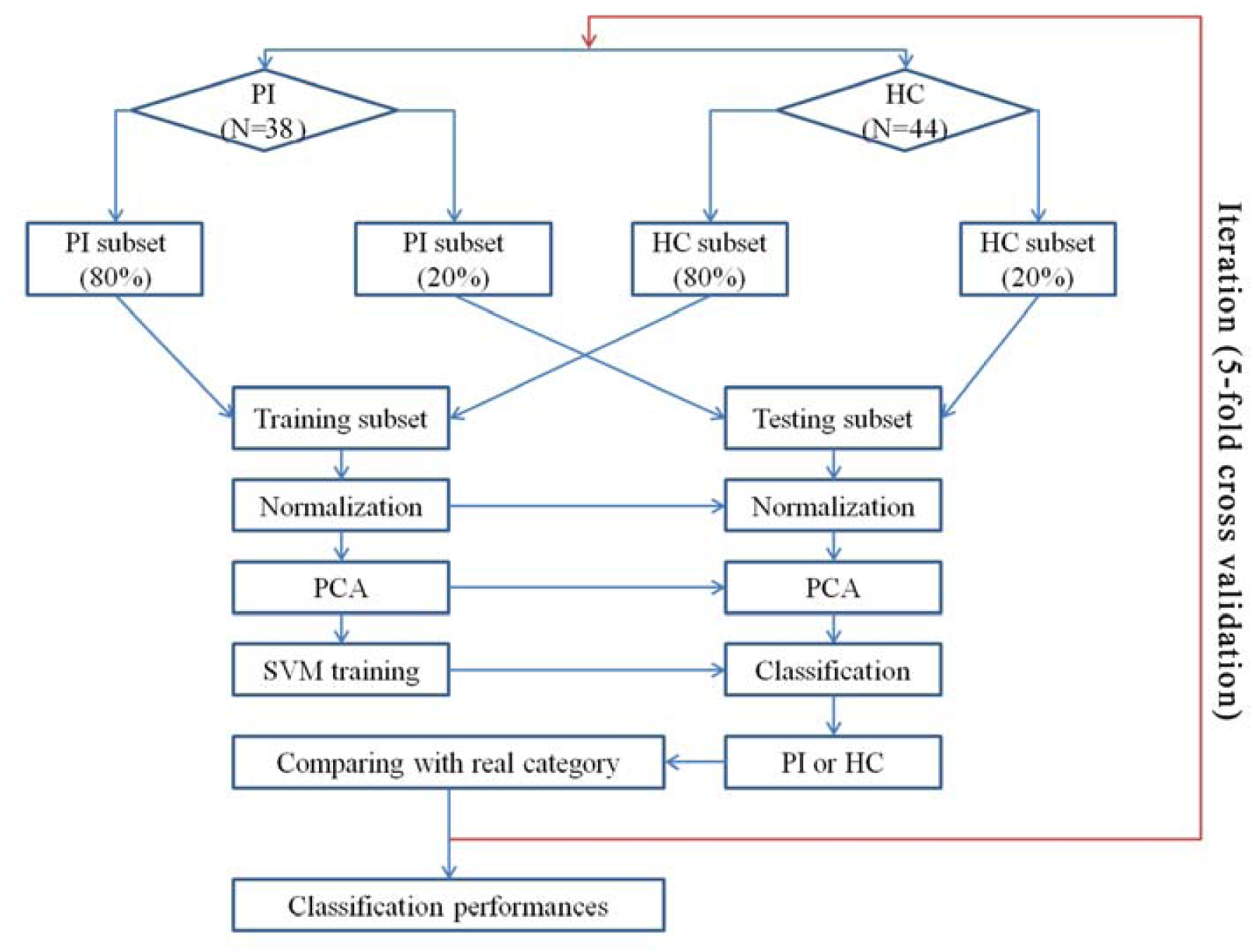
Flow chart overview illustrating the MVPA. MVPA: multivariate pattern analysis.

**Figure 2.**
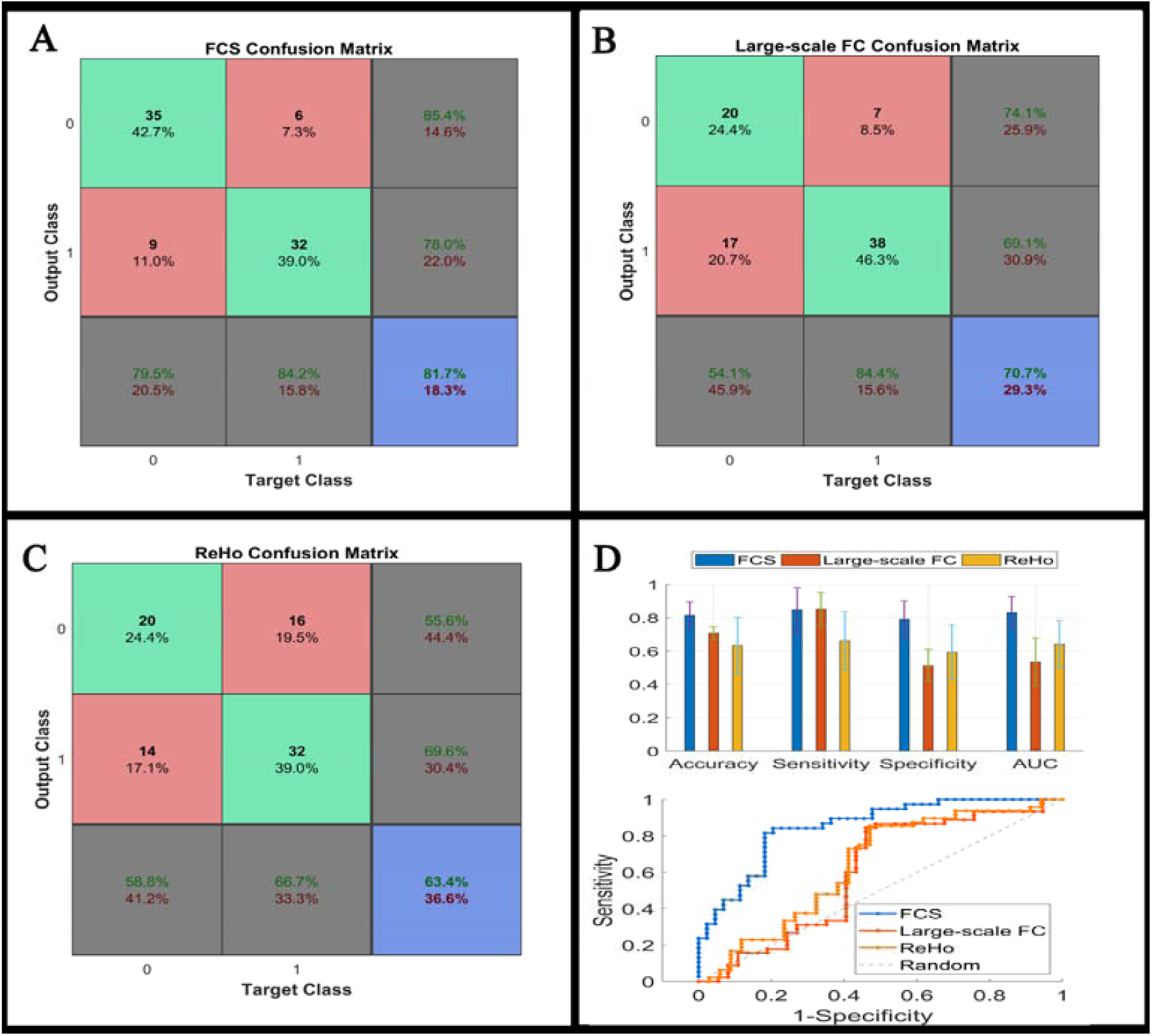
Confusion matrix (A-C), classification performances (the upper part of D) and ROC (the lower part of D) of linear SVM classifier using the three types of the functional connectivity features. Please note the subtle numerical differences between the confusion matrix and bar graph are the result of rounding. ROC: receiver operating characteristic; SVM: support vector machine.

### Statistical Analysis

Demographic and scale data of all participants were analyzed using SPSS (version 20; SPSS, Chicago, III). Differences in age, education level, Insomnia Severity Index (ISI), Pittsburgh Sleep Quality Index (PSQI), Self-rating Anxiety Scale (SAS), and Self-rating Depression Scale scores (SDS) between PI patients and healthy controls were compared using Wilcoxon rank sum tests. Differences associated with age were assessed using chi-squared tests.

Nonparametric permutation testing was used to estimate the statistical significance of the averaged classification performances by determining whether these performances exceeded chance levels. The class labels (i.e., PI patients vs. HC) of the training data were randomly permuted 1,000 times prior to training, and repeated the entire 5-fold cross-validation procedure. The P value of the permutation test was defined as: P = (N_exceed_+1)/ (N_permutation_+1). Where N_exceed_ represents the number of times the permuted performance exceeded the one obtained for the true labels. N_permutation_ represents the times of permutation.

Because of the unfavorable classification performance of the large-scale FC and ReHo (please see Figure 2), we only performed the permutation test on FCS. We additionally analyzed the between-group differences of these three FC metrics using traditional two-sample t-test, with age, sex and years of education as covariates. Since the focus of this study is FCS, correlation analysis was conducted to determine whether FCS was correlated with clinical characteristics in the PI group.

## Results

### Demographic and Scale Data

As shown in Table 1, the PI patients and the controls showed no significant differences in age (P = 0.74), sex (P = 0.10), and education level (P = 0.19). However, PI patients had higher ISI, PSQI, SAS, and SDS scores compared to HC (all P < 0.001).

**Table 1.** Demographic and Scale Data of All Study Participants

| **Variables** | **PI Group**<br>**(n=38)** | **HC Group**<br>**(n=44)** | ***P* Value** |
| --- | --- | --- | --- |
| Gender (M/F) | 16/22 | 11/33 | 0.10<sup>*</sup> |
| Age (y) | 40.61 ± 9.43 | 39.91 ± 9.43 | 0.74<sup>#</sup> |
| Duration (mo) | 40.31 ± 40.09 | N/A | N/A |
| Education (y) | 7.50 ± 3.54 | 8.45 ± 4.31 | 0.19<sup>#</sup> |
| ISI | 19.32 ± 3.09 | 5.43 ± 2.46 | < 0.001<sup>#</sup> |
| PSQI | 12.45 ± 3.09 | 5.77 ± 3.15 | < 0.001<sup>#</sup> |
| SAS | 50.29 ± 11.29 | 39.73 ± 5.68 | < 0.001<sup>#</sup> |
| SDS | 55.21 ± 9.57 | 40.39 ± 2.54 | < 0.001<sup>#</sup> |

Note.—Unless otherwise noted, data are mean ± standard deviation.
\*—The *P* value was obtained using the chi-square test.
#—The *P* value was obtained using the Wilcoxon rank sum tests.

**Table 2.** Top One Percent of the Region Weights from SVM Classifier Using the Functional Connectivity Strength

| Brain regions | MNI coordinates (Peak) |  |  | Cluster size<br>(Voxels) | weight<br>(Peak) |
| --- | --- | --- | --- | --- | --- |
|  | X | Y | Z |  |  |
| Fronto-insular cortex (R) | 39 | 3 | -6 | 122 | 0.03 |
| Precuneus (L) | -3 | -66 | 60 | 114 | -0.03 |
| Middle frontal gyrus (L) | -33 | 21 | 51 | 212 | 0.05 |

L, Left; R, Right.

### Classification Performances

Figure 2 shows the confusion matrix and classification performances of the 3 modalities. FCS reached 81.5±9.0% for accuracy, 84.9±14.7% for sensitivity, 79.1±12.3% for specificity, and 83.0±10.8% for AUC (all P < 0.001). However, several performances for large-scale FC and ReHo were around 50%, i.e. the chance level. Consequently, the focus of our study was only on FCS. Please note the subtle numerical differences between the confusion matrix and bar graph, which were the result of rounding.

### Classification Weight Maps

Figure 3 and Figure S1 show the top one percent of classification weight maps from linear SVM classifier using the FCS (cluster size threshold = 100). Right fronto-insular cortex, left precuneus and left middle frontal gyrus contributed high weight to the classifier.

**Figure 3.**
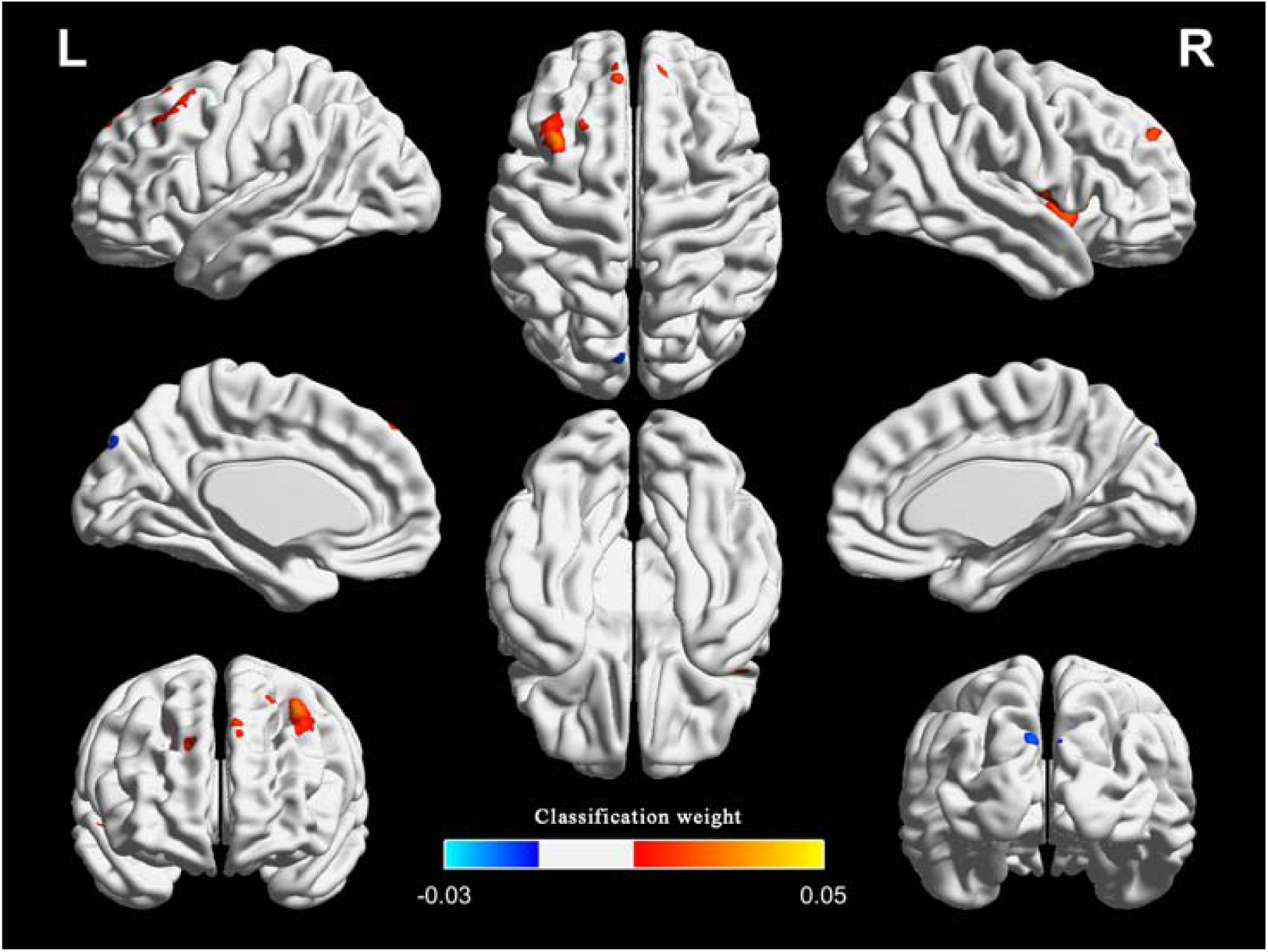
The top one percent of classification weight maps from the linear SVM classifier using the FCS as feature (cluster size threshold = 100). FCS, functional connectivity strength. FCS: functional connectivity strength, PI: primary insomnia, HC: healthy controls.

### MVPA Results of More Rigorous Inclusion Criteria

Considering that previous research used more rigorous inclusion criteria: duration > 6 month, total sleep time ≤ 6.5 h and either sleep onset latency (SOL) > 45 min or WASO > 45 min or SOL + WASO > 60 min ^28,29^, we also adopted the additional specific severity criteria to the patients group and repeated the MVPA for FCS (number of patients=22; duration=61.7±69.2; total sleep time=326.8±35.0; SOL=46.8 ±28.8; WASO=95.0±52.3). Results showed that the classification performances were 76.6±9.3% for accuracy, 76.3±9.5% for sensitivity, 76.9±13.7% for specificity, and 86.0±7.0% for AUC. The right fronto-insular cortex, left middle frontal gyrus and bilateral superior frontal gyrus had relatively high classification weights (right fronto-insular cortex and left middle frontal gyrus were the repeated regions in the two analyses). We reported the results that adopted the specific criteria in the supplementary material (Figure S2).

In addition, as to the healthy controls, substantial studies reported that PSQI total score□<□5 was defined to the healthy controls ^30-32^. In order to minimize the influence of PSQI on the results, we use linear regression method to remove the covariate PSQI. Then, we repeated the MVPA for FCS. Results showed that the classification performances were 82.9±5.9% for accuracy, 84.7±13.4% for sensitivity, 80.7±13.1% for specificity, and 90.7±5.1% for AUC. The right fronto-insular cortex and left middle frontal gyrus had relatively high classification weights (these two regions were all the repeated regions in the two analyses). We reported the results in the supplementary material (Figure S3).

### Between-group Differences and Correlation Analysis

Figure 4, Figure S4 and Table 3 illustrate the regions showing between-group differences in FCS maps (Alphasim correction for multiple comparisons of P < 0.05 combined with single voxel P < 0.01). The estimated Gaussian filter width (FWHM, in mm) were [7.161, 7.834, and 7.771]. The number of Monte Carlo simulations was 1000. Compared with HC, PI patients showed increased FCS in right fronto-insular cortex and left middle frontal gyrus, while decreased in the right head of the caudate nucleus. It is worth noting that the right fronto-insular cortex and left middle frontal gyrus also showed high classification weights.

**Table 3.** Between-group Differences (PI-HC) for Functional Connectivity Strength

| Brain Regions | MNI Coordinates (Peak) |  |  | Cluster Size<br>(Voxels) | *T* |
| --- | --- | --- | --- | --- | --- |
|  | X | Y | Z |  | Values<br>(Peak) |
| Fronto-insular cortex (R) | 36 | 3 | -6 | 100 | 5.40 |
| Head of caudate nucleus (R) | 9 | 12 | 9 | 70 | -4.85 |
| Middle frontal gyrus (L) | -33 | 21 | 51 | 64 | 4.58 |

L, Left; R, Right.
The significance level was set to $P < 0.01$ at the voxel level, with Alphasim corrections for multiple comparisons of $P < 0.05$ .

**Figure 4.**
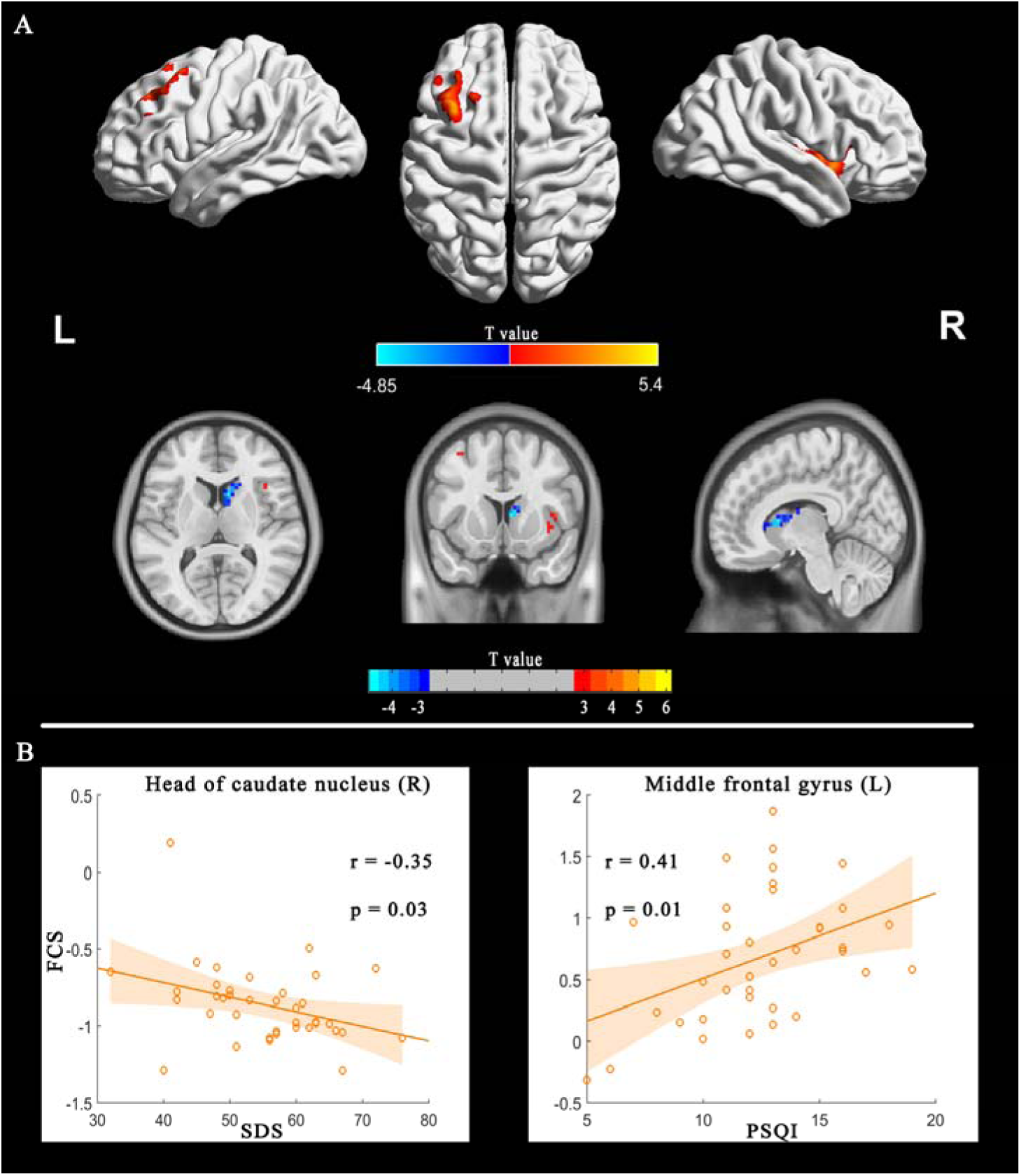
FCS differences between PI patients and HC (PI-HC). The threshold was P < 0.01 at the voxel level, with Alphasim corrections for multiple comparisons of P < 0.05. The color bar represents the t value (A). Correlation between FCS and sleep scales in the PI group.

In addition, we found that patients with primary insomnia showed increased ReHo in bilateral anterior cingulate gyrus, left precentral gyrus and superior frontal gyrus (Alphasim correction for multiple comparisons of P < 0.05 combined with single voxel P < 0.01). The estimated Gaussian filter width (FWHM, in mm) were [7.276, 8.014, and 7.841]. The number of Monte Carlo simulations was 1000. However, larger scale FC showed no between-group difference (FDR q < 0.05). We have added the between-group difference in ReHo to supplementary material (Figure S5).

Correlation analyses showed the FCS in the left middle frontal gyrus and right head of the caudate nucleus were correlated with SDS and PSQI respectively (Figure 4.B).

## Discussion

To our knowledge, this is the first study to employ MVPA for the automatic classification of patients with PI using three types of FC features. In the current study, we investigated whether or not the three types of FC metrics could be used as biomarkers to define PI. Specifically, the classification performances of FCS were all approximately equal to or more than 80% for diagnosing PI patients. The right fronto-insular cortex and left middle frontal gyrus not only had higher classification weights, but also were the repeated regions with those of between-group comparison. In addition, correlation analysis showed that FCS in left middle frontal gyrus and head of right caudate nucleus were correlated with PSQI and SDS respectively.

Convergent findings based on functional MR imaging support that spontaneous neural activity or FC in the insular cortex, prefrontal cortex and precuneus were disrupted in patients with insomnia or subjects with insomnia symptoms ^5-11^. In line with previous findings, our study firstly showed that these regions also have high classification weights.

Intriguingly, right fronto-insular cortex and left middle frontal gyrus not only had high classification weights, but also showed differences between groups. Right fronto-insular cortex is a key node of the salience network, and is implicated in arousal and insomnia ^6,33^. Michael C. Chen et al. demonstrated that the anterior insular cortex had greater involvement with the salience network, which indicated that the region was involved in hyperarousal in insomnia, and may be an important target for novel therapies for PI ^6^. Our findings that the right fronto-insular cortex had high classification weight and increased FCS might offer further confirmation.

However, our findings that increased FCS in the left middle frontal gyrus were not consistent with previous studies. Reduced metabolism, activation or spontaneous neural activities in the prefrontal cortex are the general findings ^3,9,20,34^. One explanation might be that increased FCS, a manifestation of increased interaction between a given region and other regions, was compensatory to the above reduction in the prefrontal cortex. Future researches need to verify this hypothesis.

In addition, we also found decreased FCS in the head of the caudate nucleus, which also negatively correlated with PSQI. Previous studies have established that the caudate is involved in the most consistently reported abnormalities for insomnia, i.e., hyper-arousal, sleep problems and deficits in working memory, episodic memory and problem solving ^13^. Furthermore, stimulating the caudate could reduce excitability of the human cortex ^35^. Using functional MR imaging, Diederick Stoffers et al. found that hyper-arousal, a most prominent characteristic of insomnia, was associated with reduced caudate recruitment when performing an executive task ^13^. Interestingly, our study found that the functional interaction between the head of the caudate nucleus and other brain regions was weaker at the resting state. Although FCS in the head of the caudate nucleus cannot be used to identify insomnia, decreased FCS in this region might be the underlying neurobiological substrate for hyper-arousal in insomnia.

Several limitations of the current study have to be acknowledged. First, our sample size was relatively small. Further studies using larger cohorts and multi-center imaging datasets are needed to confirm our findings. Second, we only used functional MR imaging data. The integration of structural with functional data may be a more effective method to elucidate disease factors that are shared across different modalities. Third, we only investigated the static features of the three types of FC and did not study their dynamic features. Increasing evidence has demonstrated that the functional brain connectivity have dynamic characteristics, emergent over time scales spanning milliseconds and tens of minutes. Future studies using dynamic FC are needed when performing MVPA for PI.

In summary, these limitations notwithstanding, our findings suggest that abnormal FCS in the right fronto-insular cortex and left middle frontal gyrus might serve as a potential neuromarkers for PI.

## Acknowledgements

This study was funded by the National Natural Science Foundation of China (grant no.: 81471639); the National Natural Science Foundation of China (grant no.: 81771807); the Natural Science Foundation of Guangdong Provincial (grant no.: 2015A030313723); the Science and Technology Foundation of Guangdong Province (grant no.: 2016A020215125; 2017A020215077); and the Science and Technology Foundation of Guangzhou City (grant no.: 201607010056).

## Abbreviations

PI: primary insomnia
HC: healthy controls
FC: functional connectivity
MVPA: multivariate pattern analysis
SVM: support vector machine
FCS: functional connectivity strength
ROC: receiver operating characteristic curve
AUC: area under the receiver operating characteristic curve
ISI: Insomnia Severity Index
PSQI: Pittsburgh Sleep Quality Index
SAS: Self-rating Anxiety Scale
SDS: Self-rating Depression Scale.

