## supplemental figures for "Multivariate Pattern Classification of Primary Insomnia Using Three Types of Functional Connectivity Features"

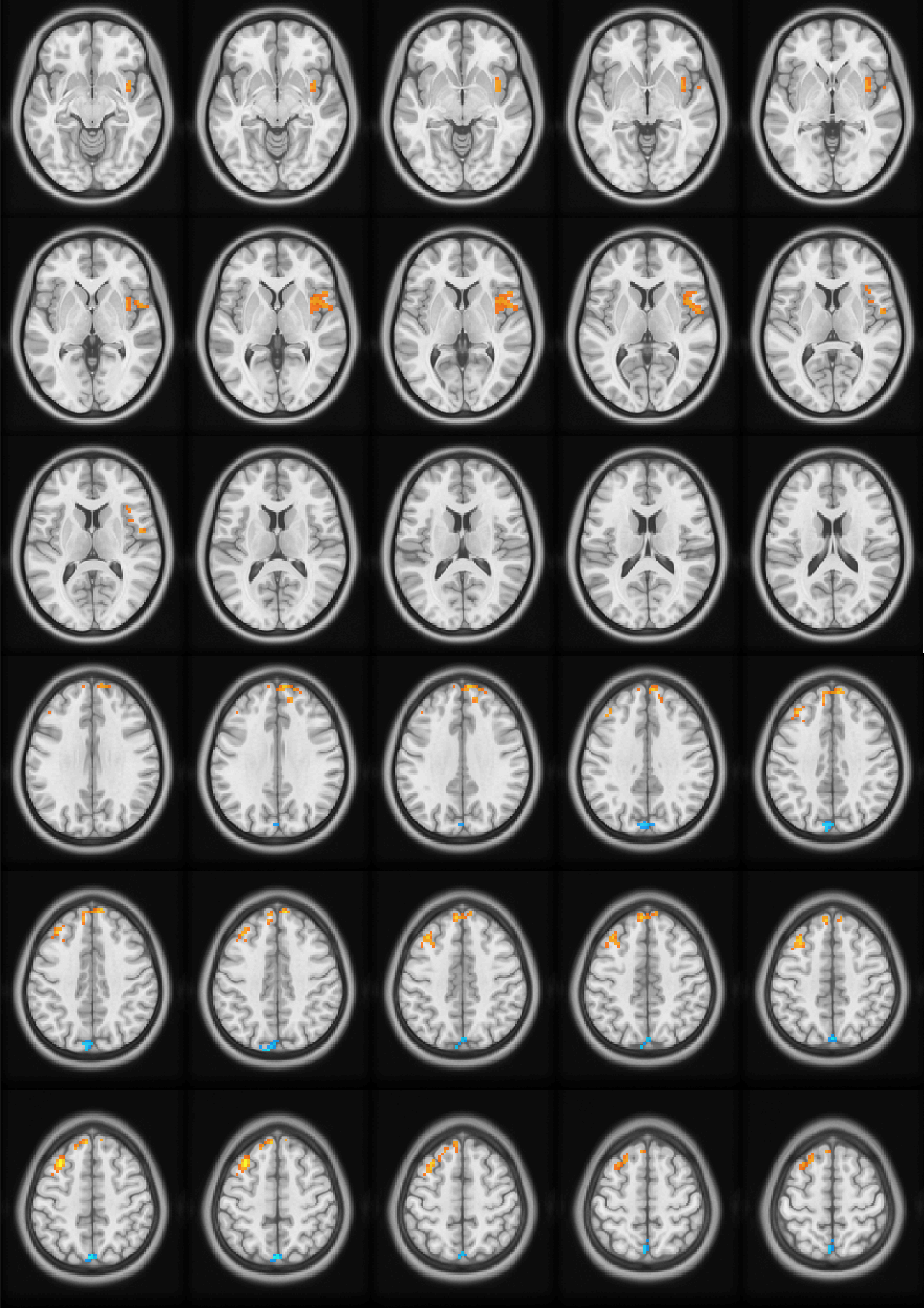

**Figure S1**. The top one percent of classification weight maps from the linear SVM classifier using the FCS as feature (cluster size threshold = 100).

FCS, functional connectivity strength.

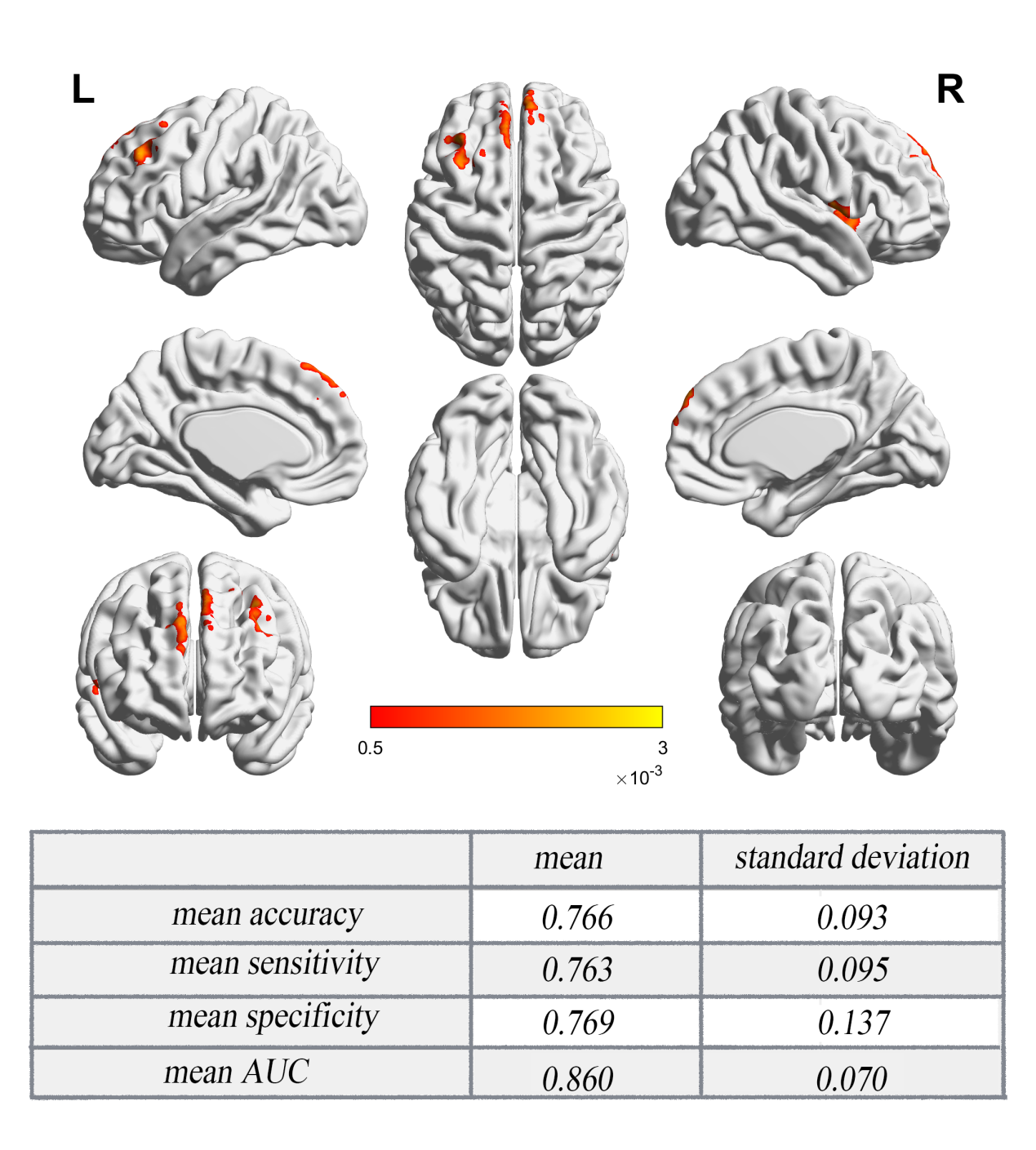

**Figure S2**. The top one percent of classification weight maps from the linear SVM classifier and classification performances using the FCS as feature using more rigorous inclusion criteria (cluster size threshold = 100).

The color bar represents the beta value.

FCS, functional connectivity strength.

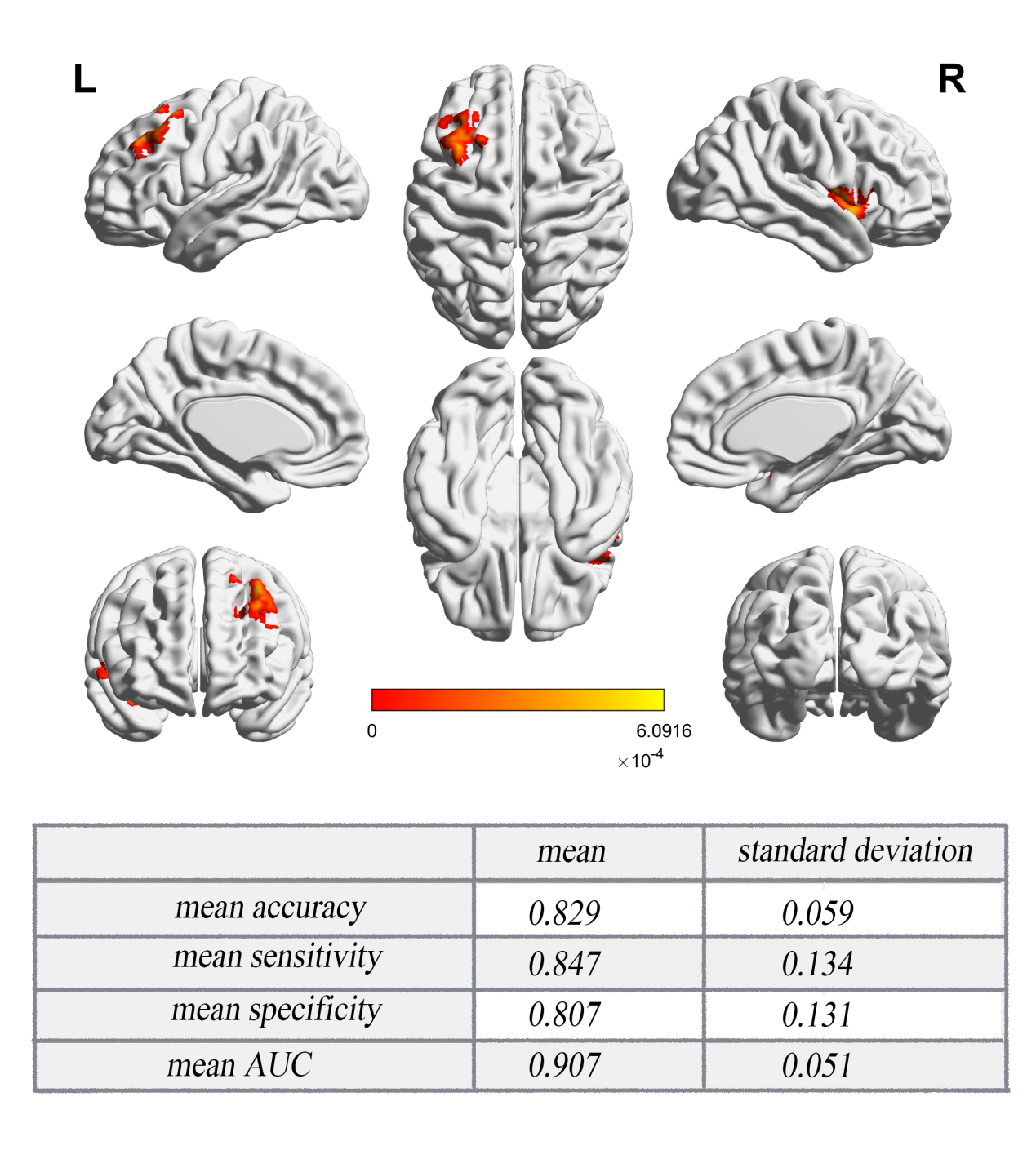

**Figure S3**. The top one percent of classification weight maps from the linear SVM classifier and classification performances using the FCS as feature when control the covariate PSQI (cluster size threshold = 100).

The color bar represents the beta value.

FCS, functional connectivity strength.

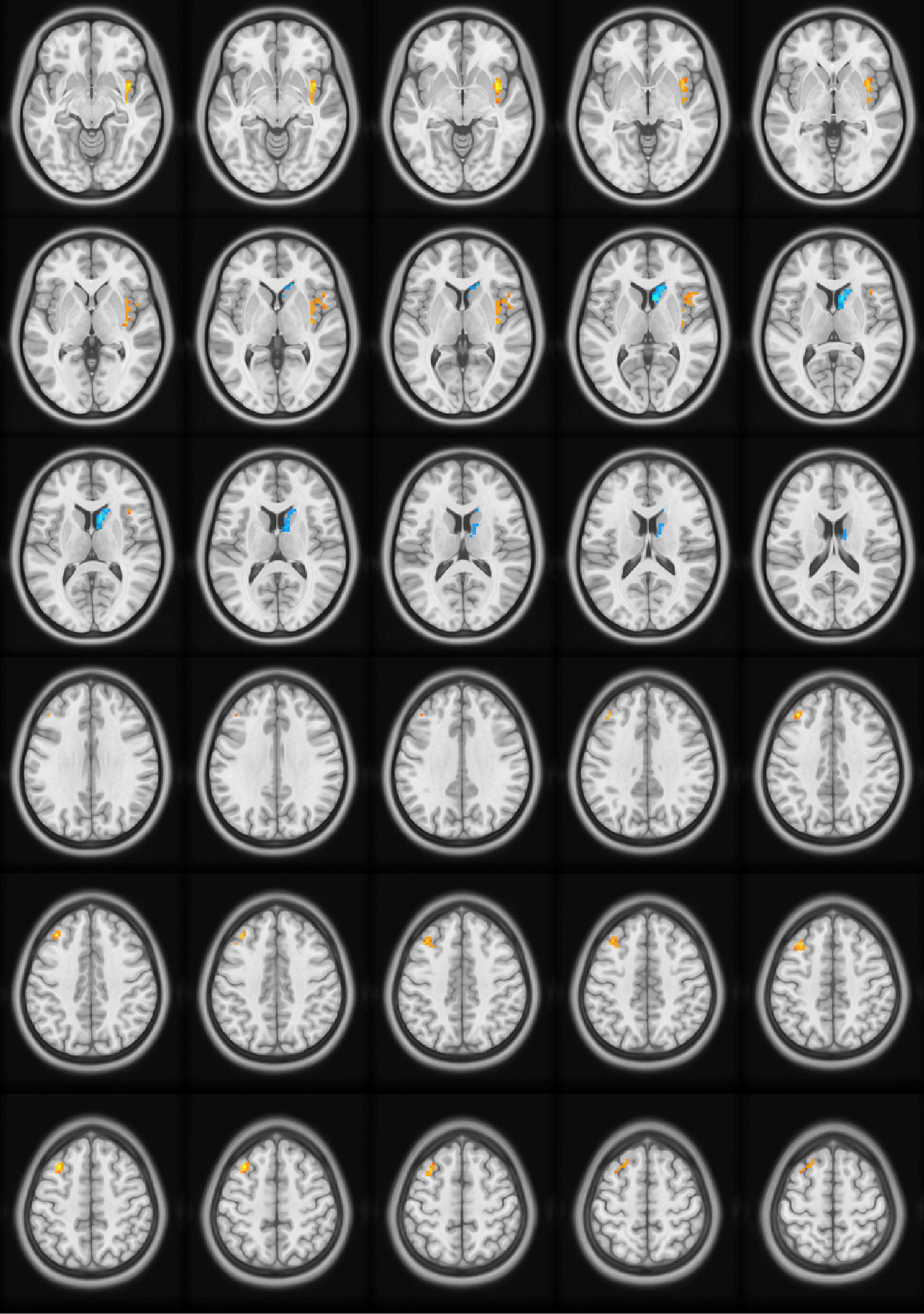

**Figure S4**: FCS differences between PI patients and HC (PI-HC). The threshold was P < 0.01 at the voxel level, with Alphasim corrections for multiple comparisons of P < 0.05.

FCS: functional connectivity strength, PI: primary insomnia, HC: healthy controls.

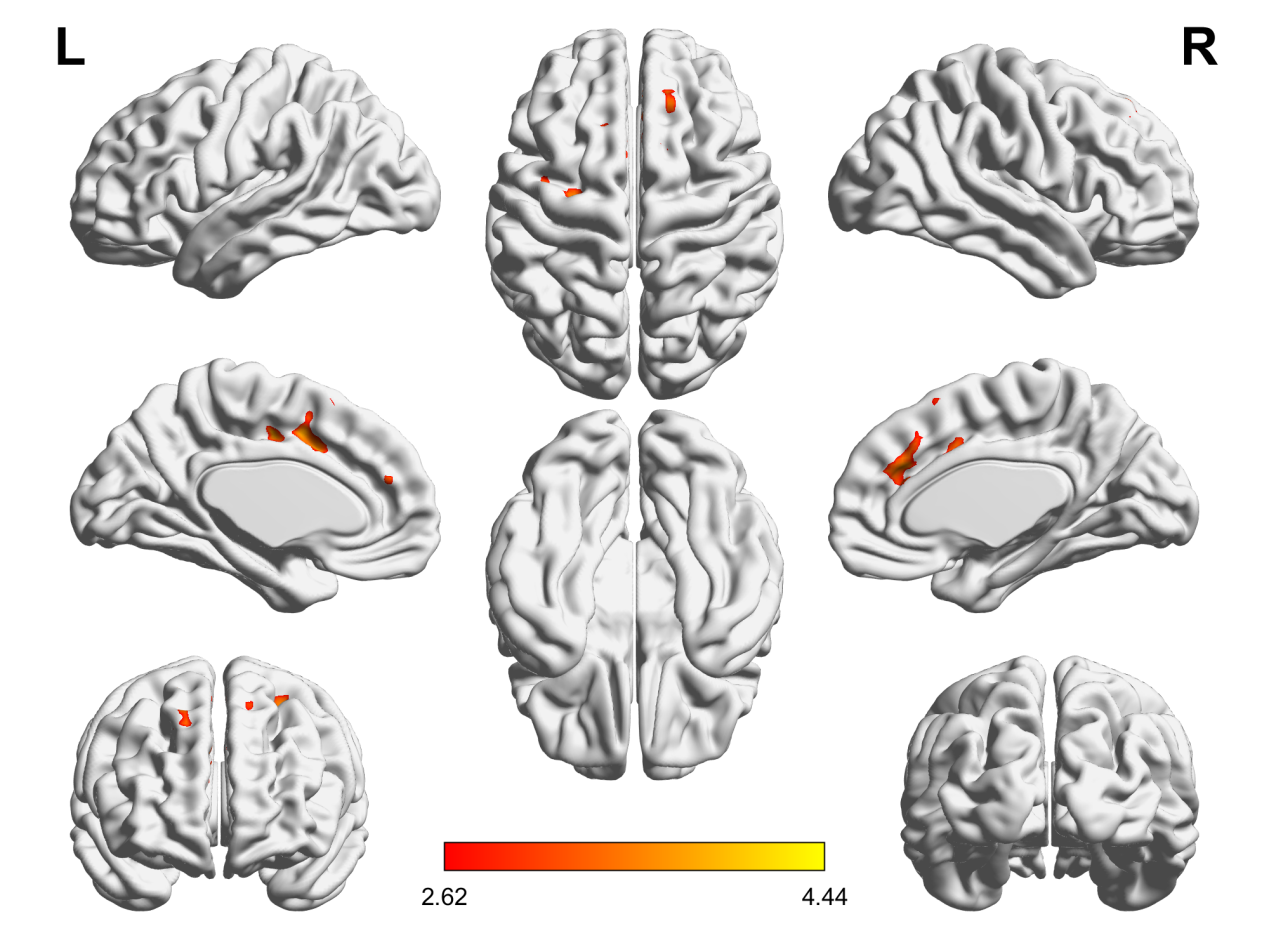

**Figure S5**: ReHo differences between PI patients and HC (PI-HC). The threshold was P < 0.01 at the voxel level, with Alphasim corrections for multiple comparisons of P < 0.05.

The color bar represents the t value.

ReHo: regional homogeneity, PI: primary insomnia, HC: healthy controls.
